# Executive control of positive and negative information and adolescent depressive symptoms: cross-sectional and longitudinal associations in a population-based cohort

**DOI:** 10.1101/548834

**Authors:** Gemma Lewis, Katherine S. Button, Rebecca M Pearson, Marcus R Munafo, Glyn Lewis

**Author notes:** Correspondence to: Dr Gemma Lewis, UCL Division of Psychiatry, 6th Floor, Maple House, 149 Tottenham Court Road, London W1T 7NF, 0203 108 7183.

## Abstract

**Background:** Large population-based studies of neuropsychological factors that characterize or precede depressive symptoms are rare. Most studies use small case-control or cross-sectional designs, which may cause selection bias and cannot test temporality. In a large UK population-based cohort we investigated cross-sectional and longitudinal associations between executive control of positive and negative information and adolescent depressive symptoms.

**Methods:** Cohort study of 2315 UK adolescents (ALSPAC) who completed an affective go/no-go task at age 18. Depressive symptoms were assessed with the Clinical Interview Schedule Revised (CIS-R) and short Mood and Feeling Questionnaire (sMFQ) at age 18, and with the sMFQ 15 months later. Analyses were linear multilevel regressions (for cross-sectional associations) and traditional linear regressions (for longitudinal associations), before and after adjustment for confounders.

**Results:** Cross-sectionally, at age 18, there was some evidence that adolescents with more depressive symptoms made more errors in executive control (after adjustments, errors increased by 0.17 of a point per 1 SD increase in sMFQ score, 95% CI 0.08 to 0.25). However, this cross-sectional association was not observed for the CIS-R (.03, 95% CI -.06 to .12). There was no evidence of a difference in executive control errors according to valence. Longitudinally, there was no evidence that reduced executive control was associated with future depressive symptoms.

**Conclusions:** Executive control of positive and negative information does not appear to be a marker of current or future depressive symptoms in adolescents and would therefore not be a useful target in interventions to prevent adolescent depression. According to our evidence, the affective go/no-go task is also not a good candidate for future neuroimaging studies of adolescent depression.

## Introduction

Depression is the leading cause of disease burden worldwide, with no generally accepted methods of prevention (Vos, 2016). Poor cognitive functioning is an important cause of the disability associated with depressive illness (Roiser & Sahakian, 2013) and may be a cause of depression, rather than just a consequence. However, the neuropsychological processes underlying cognitive dysfunction in depression are poorly understood. One theory of depression suggests there is reduced connectivity between neural circuits involved in executive control (such as the prefrontal cortex) and neural circuits that respond to positive and negative emotional information (such as the ventral striatum) (Furman, Hamilton, & Gotlib, 2011; Hamilton, Chen, Thomason, Schwartz, & Gotlib, 2011; Treadway & Pizzagalli, 2014). The precise mechanisms are poorly understood but may involve disrupted translation of reward sensitivity into reward seeking behaviours (Furman et al., 2011). If this were true, reduced top down inhibition by the prefrontal cortex would be a risk factor for later depression in longitudinal neuroimaging studies.

This question has proved difficult to study. Neuroimaging studies often use small convenience samples, which may lack statistical power and increase the possibility of an unreliable finding (Button et al., 2013; LeWinn, Sheridan, Keyes, Hamilton, & McLaughlin, 2017). Small samples can also make findings more difficult to generalise. This is particularly true for neuroimaging studies which, even when large, have strict exclusion criteria (e.g people with tattoos and claustrophobia). Sample composition probably introduces selection bias and contributes to the poor reproducibility of many neuroimaging findings (LeWinn et al., 2017). Finally, the reliability of many neuroimaging measures, including those of executive functioning and emotional processing, is very low (Nord, Gray, Charpentier, Robinson, & Roiser, 2017; Plichta et al., 2012).

An alternate strategy is to investigate behavioural performance on neuropsychological tasks, which can be embedded in larger epidemiological studies that allow more robust conclusions about any cross sectional or longitudinal associations. If performance on a task is associated with depressive symptoms, the neural mechanisms underlying the behavior can then be investigated using neuroimaging.

Executive control of positive and negative information can be measured using the affective go/no-go task, which has been shown to engage several regions of the prefrontal cortex (Elliott, Rubinsztein, Sahakian, & Dolan, 2002). The affective go/no-go task has been used in small case-control studies of adults and adolescents, to identify abnormalities in the executive control of positive and negative information that might characterise depressive illness (Erickson et al., 2005; Kyte, Goodyer, & Sahakian, 2005; Maalouf et al., 2012; Murphy et al., 1999). However, findings from these studies are very inconsistent (Supplementary Table 1). One limitation of case-control studies is that they are more prone to selection biases than cross-sectional or cohort studies because of the difficulty in selecting controls from the same population as the cases (Rothman, Greenland, & Lash, 2013). It is also impossible using a case-control design to tell the direction of any association. We are only aware of two cohort studies of this question (Kilford et al., 2015; Owens et al., 2012) and their findings are inconsistent (Supplementary Table 1). Both studies were of small samples (less than 263 adolescents) selected for high-risk of depression, which may have reduced statistical power or introduced selection bias.

In this study we used a large population-based cohort of UK adolescents to compare cross-sectional and longitudinal associations between executive control of positive and negative information and depressive symptoms. Our aim was to distinguish between abnormalities in executive control that result from, or are concurrent with, depressive symptoms and those that are associated with future risk. The influence of valence on the association between executive control and depressive symptoms is inconclusive, so we tested whether adolescents with depressive symptoms would show worse executive control in response to positive than negative information.

## Methods

### Participants

The Avon Longitudinal Study of Parents and Children (ALSPAC) is an ongoing population-based birth cohort examining a wide range of influences on health and development (Boyd et al., 2013; Fraser et al., 2013). All pregnant women living in the former county of Avon in Bristol, South West England (UK), with an estimated delivery date between April 01 1991 and December 31 1992 were invited to participate. The core enrolled sample consisted of 14541 women (an estimated 85-90% of those eligible) and 13154 fathers/partners. An additional 713 children were enrolled during phases 2 and 3 of the study. The total sample size for analyses was 15 247 pregnancies with 15 458 fetuses. Of this total sample, 14 775 (95.6%) were live births and 14 701 infants (95.1%) were alive at 1 year of age. Mothers, fathers and offspring have regularly provided data, either through postal questionnaires or in research clinics. Further information about ALSPAC is available on the study website (www.bristol.ac.uk/alspac), which includes a fully searchable data dictionary (www.bris.ac.uk/alspac/researchers/data-access/data-dictionary). Ethical approval for the study was obtained from the ALSPAC Ethics and Law Committee and the Local Research Ethics Committees. In this study we used data from core singleton offspring who completed the affective go/no-go task at a research clinic when they were 18 years old. All participants provided written informed consent.

### Measures

#### Depressive symptoms

We used two measures of depressive symptoms that were available at age 18, the Clinical Interview Schedule Revised (CIS-R) and the short Mood and Feelings Questionnaire (sMFQ). The CIS-R is a self-administered computerised clinical assessment, which assesses symptoms of common mental disorder during the past week. The CISR can be used to generate ICD-10 diagnoses of depression, a total depressive symptom score, and a total score that reflects the overall severity of common mental disorder psychopathology (Lewis, Pelosi, Araya, & Dunn, 1992). The depression score is calculated by summing scores for the following CIS-R items: depression, depressive ideas, fatigue, poor concentration and sleep disturbance. Possible scores range from 0 to 21, higher scores indicating more severe depressive symptoms. In this study we used the depressive symptom score as our primary outcome. Symptoms of depression and anxiety frequently co-exist and many people meet criteria for more than one common mental disorder. As a supplementary analysis we also tested associations with the CIS-R total score which measures symptoms of six types of common mental disorder (depression, generalised anxiety disorder, panic disorder, phobias and obsessive compulsive disorder).

The short MFQ was administered at age 18 and also 15 months later, so was used in longitudinal as well as cross-sectional analyses. The short MFQ is a 13-item self-report measure of the severity of DSM-IV depressive symptoms in the past two weeks. Possible scores range from 0 to 26, higher scores indicating more severe depressive symptoms (Turner, Joinson, Peters, Wiles, & Lewis, 2014).

#### Affective go/no-go task

Participants completed the task at a research clinic when they were an average age of 17.5 years (N = 2484). Single words are flashed onto the centre of a computer screen, and are either positive (hopeful, serene) or negative (glum, mistake). Each word is displayed for 300 milliseconds, with 900 milliseconds intervals. The task is split into 8 blocks (two practice and two experimental) of 18 words (nine positive and nine negative). Participants were initially instructed to press the space bar as fast as they could for positive words (‘targets’) but not negative words. After two word blocks requiring responses to positive words, the instructions change so that the space bar is to be pressed for sad words (‘shifts’). Conditions are alternated in a PPNNPPNN pattern to create shift and nonshift response blocks. Measures extracted from the affective go/no-go task include: 1) commission errors (how many times participants respond or ‘go’ to a non-target word, for example pressing the space bar in response to a negative word when positive words are targets) 2) omissions (how many times participants miss a target word, for example not pressing the space bar in response to a negative word when negative words are targets and 3) time taken to respond to target words (reaction times).

#### Potential confounders

We adjusted for variables previously shown to be associated with exposure and outcome and not on the causal pathway. These included sex, age at time of the research clinic and Intelligence Quotient (IQ) measured at the closest possible time-point to the exposure (age 15). Participants were administered the Vocabulary and Matrix Reasoning subsections of the Wechsler Abbreviated Scale of Intelligence (Wechsler, n.d.). IQ was calculated for each participant, adjusted for their age. We also adjusted for maternal education and social class as indicators of socioeconomic position. Educational qualification was coded 1 to 5, ranging from Certificate of Secondary Education (which used to be compulsory in the UK) to university degree. This was dichotomized to create a binary variable (compulsory and non-compulsory education). Social class was measured using five categories from the 1991 classification of the UK Office of Population Censuses and Surveys, dichotomized into manual and non-manual (Tilling, Macdonald-Wallis, Lawlor, Hughes, & Howe, 2014).

### Statistical analyses

#### Cross-sectional associations between positive and negative executive control and depressive symptoms at age 18

We used linear multilevel models to test whether there was a main effect of depressive symptoms, valence (positive or negative), shift condition or error type (commission or omission) on total number of errors, modelled as a continuous outcome. A random effect was included for participant, to account for clustering within individuals. This model was then adjusted for potential confounders. Residual distributions were inspected for normality. We tested separate models for the MFQ and CIS-R depressive and total score measures.

To test whether depressive symptoms were associated with more omission errors for positive than negative words, and fewer commission errors for positive than negative words, we calculated a three-way interaction between depressive symptoms, error type, and valence.

We also tested whether these associations were influenced by shift condition. In a separate model, we tested a four-way interaction between depressive symptoms, error type, valence, and shift condition.

#### Longitudinal associations between positive and negative executive control at age 18 and depressive symptoms 15 months later

MFQ scores 15 months after the affective go/no-go task were modelled as a continuous outcome, using linear regression. Exposure variables were positive commission errors, negative commission errors, positive omissions and sad omissions (all continuous). First we calculated univariable associations with each of these exposure variables. Next we tested a multivariable model that included all four exposures. We then adjusted this multivariable model for potential confounders, including MFQ scores at baseline.

#### Reaction times and depressive symptoms

Since some studies report associations between reaction times to respond to target words and depressive symptoms, we also explored these associations. Cross-sectionally, reaction times were modelled as a continuous outcome using linear mixed models. Depressive symptoms, valence and shift condition were exposure variables. This model was then adjusted for potential confounders. We tested a three-way interaction between depressive symptoms, valence and shift condition. Longitudinal models were examined when there was any evidence of cross-sectional associations.

#### Missing data

We used multiple imputation by chained equations (MICE) to account for missing data, because complete case analyses can introduce bias when data are not missing completely at random (Sterne et al., 2009). We assumed missingness was dependent on observed data (missing at random), and imputed 50 datasets. To predict missing data, we used all variables selected for analysis models and a number of auxiliary variables including MFQs from age 11 to 16. We imputed missing confounder and outcome data for the core singleton sample who had complete data on the affective go/no-go task and at least one prior MFQ (N = 2,315), to improve prediction of missing depression data (affective go/no-go task data were not imputed). As sensitivity analyses, we also report analyses on complete case samples.

## Results

### Descriptive statistics

The affective go/no-go task was completed by 2484 of 5215 (48%) adolescents who attended the age 18 ALSPAC clinic, 2328 of whom were members of the core singleton ALSPAC sample. Of these, 2323 (94%) completed the CIS-R and 2204 (89%) the MFQ at the same clinic. CIS-R depression diagnoses were available for 2319 (93%). Once all potential confounders were included, 1,267 adolescents (54% of the core sample who completed the task at the age 18 clinic) had complete data cross-sectionally. Of these, 743 (32% of the core sample who completed the task at the age 18 clinic) also had longitudinal data on the MFQ 15 months later.

We classed as our ‘analytic sample’ (which was later imputed), adolescents in the core sample who completed the affective go/no-go task and had at least one sMFQ in the course of the study (N = 2315). Demographic characteristics for the analytic sample are presented in Supplementary Table 2, compared to the rest of the core singleton ALSPAC cohort. Adolescents with missing data were more likely to be male, from lower social classes, have a lower IQ, and meet diagnostic criteria for depression. Characteristics of the analytic sample according to errors made on the task, are shown in Table 1.

**Table 1.** Characteristics of the analytic sample (N = 2315), according to errors made on the AGNG task split at the median.

| Characteristic | Commission errors |  | Omissions |  |
| --- | --- | --- | --- | --- |
|  | Below Median (%) | Above Median (%) | Below median (%) | Above Median (%) |
| Female (n=1307) | 60.7 | 51.4 | 56.9 | 56.0 |
| Lowest maternal education, O Level or less (n=1203) | 50.3 | 57.6 | 46.1 | 61.6 |
| Lowest maternal social class (n=137) | 17.2 | 19.5 | 15.8 | 21.1 |
| Offspring depression at age 18 (n=347)* | 6.6 | 6.0 | 7.0 | 5.7 |
| Maternal age at birth, mean (SD) | 29.5 (4.5) | 29.1 (4.6) | 29.7 (4.5) | 29.0 (4.6) |
| Offspring IQ score, mean (SD) | 94.0 (12.4) | 90.2 (12.4) | 95.5 (12.1) | 88.8 (12.1) |
| Offspring MFQ score at 18, mean (SD) | 5.9 (5.0) | 6.8 (5.5) | 6.0 (5.1) | 6.7 (5.4) |
| Offspring CIS-R score at 18, mean (SD) | 2.9 (3.7) | 2.8 (3.7) | 2.9 (3.7) | 2.8 (3.7) |

\*According to ICD-10 criteria assessed with the CIS-R.

Descriptive data for performance on the affective go/no-go task in the analytic sample overall and according to depression diagnoses on the CIS-R are shown in Table 2, and according to the clinical cut-off on the sMFQ (Table 3). Overall, adolescents made more positive than negative commission errors (Table 2). However they also made more positive than negative omissions (Table 2). The correlation between commission errors and omissions was r = 0.20, p <.0001. Adolescents were also faster to respond to positive than negative target words (Table 2). No differences were observed for CIS-R depression diagnoses (Table 2). Those who exceeded the clinical cut-off score on the MFQ made more commission errors and omissions, in response to both positive and negative words, in both shift and non-shift conditions (Table 3). Evidence for a difference in positive omissions was weak (Table 3).

**Table 2.** Mean (standard deviation) scores for affective go-no-go measures in the analytic sample overall, and according to depression diagnosed using the CIS-R at age 18 (N = 2315). Comparisons are between groups with and without depression.

| Affective go-no/go measures | Overall<br>(N=2315) | Meeting ICD-10<br>criteria<br>(N=137, 6.3%) | Not meeting<br>ICD-10 criteria<br>(N=2030, 94%) | Mean difference (95% CI) |
| --- | --- | --- | --- | --- |
| Commission errors |  |  |  |  |
| Overall | 18.0 (9.7) | 17.1 (8.5) | 18.0 (9.8) | .97 (-2.65 .72) |
| Positive, shift | 5.8 (3.3) | 5.7 (3.2) | 5.8 (3.3) | -.15 (-.72 .42) |
| Positive, no shift | 4.3 (3.1) | 4.0 (2.6) | 4.3 (3.2) | -.32 (-.86 .22) |
| Negative, shift | 4.2 (2.9) | 3.8 (2.5) | 4.2 (2.9) | -.34 (-.84 .16) |
| Negative, no shift | 3.7 (2.8) | 3.6 (2.3) | 3.7 (2.8) | -.16 (-.63 .32) |
| Omissions |  |  |  |  |
| Overall | 14.0 (11.0) | 13.7 (11.5) | 14.0 (11.0) | -.34 (-2.26 1.57) |
| Positive, shift | 5.3 (4.0) | 5.4 (4.0) | 5.3 (4.0) | .15 (-.54 .84) |
| Positive, no shift | 3.3 (3.5) | 2.9 (3.3) | 3.3 (3.6) | -.35 (-.97 .26) |
| Negative, shift | 2.8 (3.0) | 2.7 (3.1) | 2.8 (3.0) | -.08 (-.60 .44) |
| Negative, no shift | 2.7 (2.9) | 2.6 (3.0) | 2.7 (2.9) | -.06 (-.56 .44) |
| Reaction times |  |  |  |  |
| Positive targets, shift | 504.7 (81.6) | 500.5 (77.7) | 504.5 (82.1) | -3.97 (-18.12 10.18) |
| Positive targets, no shift | 518.0 (78.6) | 519.6 (69.2) | 517.4 (79.1) | 2.19 (-11.40 15.78) |
| Negative targets, shift | 517.5 (72.9) | 522.2 (59.0) | 516.7 (73.9) | 5.48 (-7.17 18.13) |
| Negative targets, no shift | 524.4 (75.0) | 529.12 (62.17) | 523.4 (75.3) | 6.06 (-6.98 19.10) |

**Table 3.**
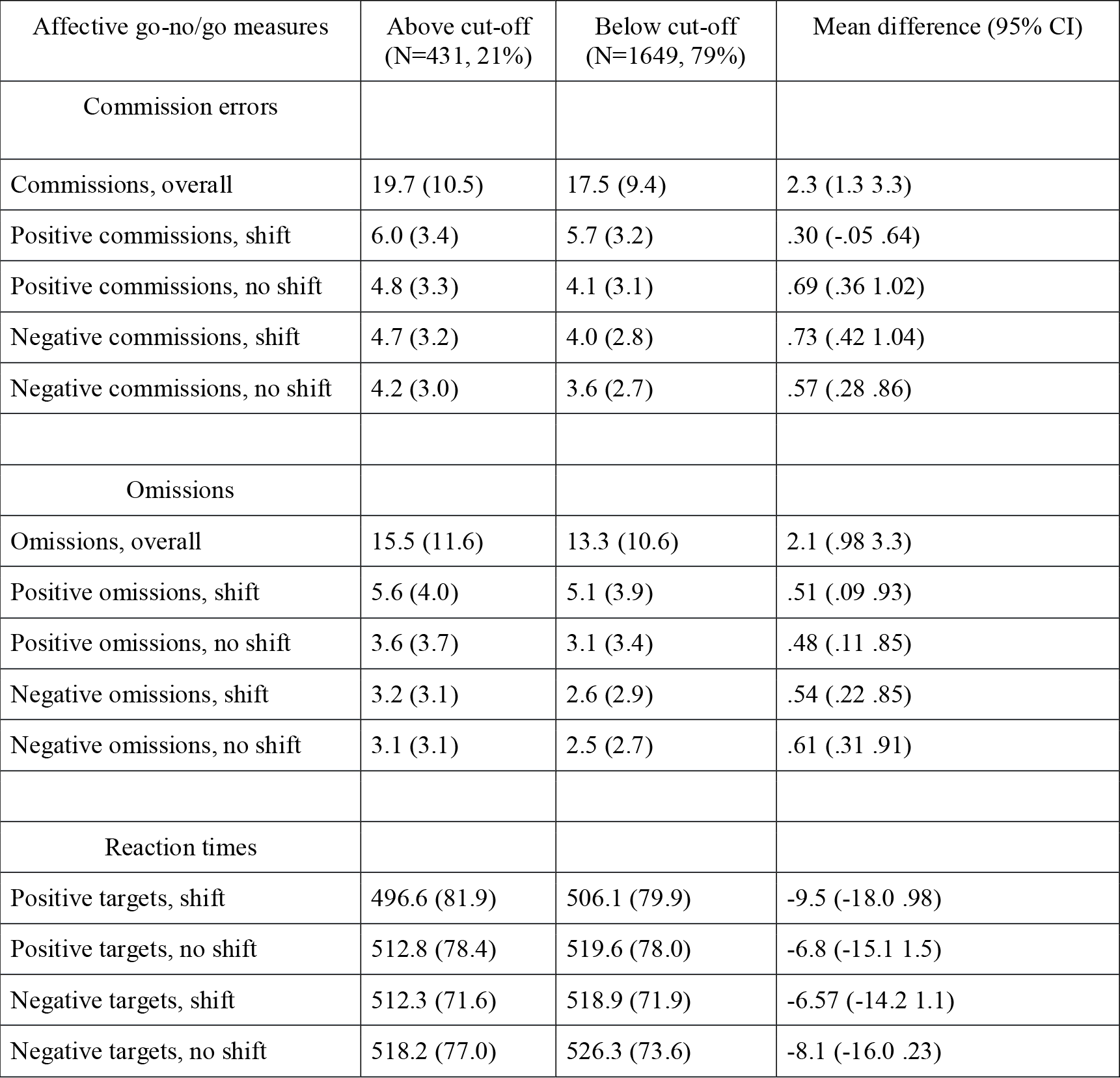
Mean (standard deviation) scores for affective go-no-go measures, according to depression (MFQ>=11) at age 18, in the analytic sample (N = 2315). Comparisons are between groups with and without depressive symptoms.

| Affective go-no/go measures | Above cut-off<br>(N=431, 21%) | Below cut-off<br>(N=1649, 79%) | Mean difference (95% CI) |
| --- | --- | --- | --- |
| Commission errors |  |  |  |
| Commissions, overall | 19.7 (10.5) | 17.5 (9.4) | 2.3 (1.3 3.3) |
| Positive commissions, shift | 6.0 (3.4) | 5.7 (3.2) | .30 (-.05 .64) |
| Positive commissions, no shift | 4.8 (3.3) | 4.1 (3.1) | .69 (.36 1.02) |
| Negative commissions, shift | 4.7 (3.2) | 4.0 (2.8) | .73 (.42 1.04) |
| Negative commissions, no shift | 4.2 (3.0) | 3.6 (2.7) | .57 (.28 .86) |
| Omissions |  |  |  |
| Omissions, overall | 15.5 (11.6) | 13.3 (10.6) | 2.1 (.98 3.3) |
| Positive omissions, shift | 5.6 (4.0) | 5.1 (3.9) | .51 (.09 .93) |
| Positive omissions, no shift | 3.6 (3.7) | 3.1 (3.4) | .48 (.11 .85) |
| Negative omissions, shift | 3.2 (3.1) | 2.6 (2.9) | .54 (.22 .85) |
| Negative omissions, no shift | 3.1 (3.1) | 2.5 (2.7) | .61 (.31 .91) |
| Reaction times |  |  |  |
| Positive targets, shift | 496.6 (81.9) | 506.1 (79.9) | -9.5 (-18.0 .98) |
| Positive targets, no shift | 512.8 (78.4) | 519.6 (78.0) | -6.8 (-15.1 1.5) |
| Negative targets, shift | 512.3 (71.6) | 518.9 (71.9) | -6.57 (-14.2 1.1) |
| Negative targets, no shift | 518.2 (77.0) | 526.3 (73.6) | -8.1 (-16.0 .23) |

### Cross-sectional associations between positive and negative executive control and depressive symptoms at age 18

We found strong evidence for an association between depressive symptoms assessed with the sMFQ and errors on the AGNG (Table 4). For every one standard deviation (SD) increase in depressive symptoms assessed using the sMFQ (5-points), total errors increased by .17 of a point (95% CI .09 to .26). This association was observed irrespective of error type, valence, and shift condition, and remained after adjustment for confounders (.17, 95% CI .08 to .25). However when using the CIS-R depressive symptom score, there was no evidence of an association before (-.03 per SD, 95% CI -.12 to .06) or after (.03, 95% CI -.06 to .12) adjustment for confounders. There was also no evidence of this association when using the CISR total score that includes both depressive and anxiety symptoms, before (-.06 per SD, -.14 to .02) or after (.01 per SD, -.07 to .09) adjustments.

**Table 4.** Change in number of errors on the affective go/no-go task, 95% confidence intervals and p value, for a one SD change in depressive symptoms. Associations are cross-sectional.

| Exposure variable<sup>a</sup> | Errors (n=2315) |  |
| --- | --- | --- |
|  | Model 1<sup>b</sup> | Model 2: model 1 adjusted for confounders<sup>c</sup> |
| MFQ depressive symptoms | .17 (.09 to .26) .0001 | .17 (.08 to .25) .0001 |
| CIS-R depressive symptoms | -.03 (-.12 to .06) .523 | .03 (-.06 to .12) .489 |
| CIS-R total score | -.06 (-.14 to .02) .164 | .01 (-.07 to .09) .838 |

<sup>a</sup>Separate models were run for depressive symptoms assessed with the MFQ and CIS-R
<sup>b</sup>Model 1 simultaneously included depressive symptoms, error type (commission or omission), valence (positive or negative) and shift condition (yes or no).
<sup>c</sup>Confounders included in model 2 were offspring age, gender and IQ and maternal education and social class

We found no evidence that the association between depressive symptoms and errors differed according to error type (commission or omission) and valence (interaction p value .684 for the MFQ and .630 for CIS-R depression score). In a separate model, we also found no evidence of an influence of shift condition (interaction p value .360 on the MFQ and .571 on the CIS-R).

### Longitudinal associations between positive and negative executive control at age 18 and depressive symptoms 15 months later

We examined commission and omission errors overall (collapsed across valence and shift), because in cross-sectional analyses we found no evidence that adjusting for valence or shift condition influenced the association between errors and depressive symptoms. We also present commission and omission errors separately by valence (Table 5). We found no evidence of an association between commission (adjusted coefficient: .01, 95% CI -.02 to .03) or omission errors (adjusted coefficient: -.01, 95% CI -.03 to .02) and later depressive symptoms. These associations were similar when conducted separately by valence.

**Table 5.** Change in depressive symptoms, 95% confidence intervals and p value, for a one point increase in errors. Longitudinal associations.

| Exposure variable<sup>a</sup> | Depressive symptoms (n=2115) |  |  |  |
| --- | --- | --- | --- | --- |
|  | Univariable<sup>b</sup> | Bivariable<sup>c</sup> | Further adjusted for confounders<sup>d</sup> | Further adjusted for baseline depressive symptoms |
| Commissions | .02 (-.00 to .05) .100 | .02 (-.01 to .05) .145 | .03 (-.00 to .06) .051 | .01 (-.02 to .03) .656 |
| Omissions | .01 (-.01 to .04) .262 | .01 (-.01 to .04) .416 | .00 (-.02 to .03) .753 | -.01 (-.03 to .02) .628 |
| Exposure variable<sup>e</sup> | Depressive symptoms (n=2115) |  |  |  |
|  | Univariable<sup>f</sup> | Multivariable<sup>g</sup> | Further adjusted for confounders<sup>d</sup> | Further adjusted for baseline depressive symptoms |
| Happy commissions | .03 (-.02 to .08) .180 | .02 (-.05 to .08) .601 | .02 (-.04 to .09) .461 | .01 (-.05 to .07) .732 |
| Sad commissions | .04 (-.01 to .10) .109 | .03 (-.05 to .10) .479 | .03 (-.04 to .11) .352 | .00 (-.06 to .07) .960 |
| Happy omissions | .02 (-.03 to .06) .434 | -.00 (-.06 to .05) .881 | -.01 (-.07 to .05) .724 | -.02 (-.07 to .04) .545 |
| Sad omissions | .03 (-.02 to .08) .184 | .03 (-.04 to .10) .372 | .02 (-.04 to .09) .471 | .01 (-.05 to .07) .810 |

<sup>a</sup>Commission and omission errors are collapsed across valence (positive and negative)
<sup>b</sup> Separate univariable models for commission and omission errors
<sup>c</sup>Includes both commission and omission errors
<sup>d</sup>Adjusted for offspring age, gender and IQ and maternal education and social class.
<sup>e</sup>Happy and sad commission errors and happy and sad omissions (collapsed across shift)
<sup>f</sup>Separate univariable models for happy commissions, sad commissions, happy omissions, sad omissions (collapsed across shift)
<sup>g</sup>Includes happy and sad commission errors and happy and sad omissions (collapsed across shift)

### Cross-sectional associations between reaction times for hits and depressive symptoms at age 18

Associations are shown in Table 6. There was some evidence that adolescents with more severe depressive symptoms responded more quickly to target words (hits), but this was weak and attenuated further after adjustment for confounders. Reaction times for hits were faster in response to positive than negative targets, and in shift than non-shift conditions. There was no evidence of interaction between depressive symptoms and valence (p value for the MFQ .253 and CIS-R .371) or depressive symptoms and shift condition (p value for the MFQ .406 and CIS-R .838).

**Table 6.** Change in reaction times for hits, 95% confidence intervals and p value, for a one SD increase in depressive symptoms

| Exposure variable <sup>a</sup> | Reaction times (n=2315) |  |
| --- | --- | --- |
|  | Model 1 <sup>b</sup> | Model 2: model 1 adjusted for confounders <sup>c</sup> |
| MFQ depressive symptoms | -2.45 (-5.29 .38) .090 | -1.51 (-4.39 1.47) .304 |
| CIS-R depressive symptoms | -.98 (-2.00 3.96) .517 | 1.42 (-1.63 4.48) .361 |
| CISR total score | -.68 (-3.56 to 2.21) .645 | -.31 (-3.28 to 2.65) .836 |

<sup>a</sup>Separate models were run for symptoms assessed with the MFQ and CIS-R
<sup>b</sup>Model 1 simultaneously included depressive symptoms, error type (commission or omission), valence (positive or negative) and shift condition (yes or no).
<sup>c</sup>Confounders included in model 2 were offspring age, gender and IQ and maternal education and social class.

## Discussion

We found some evidence that adolescents with depressive symptoms had reduced executive control of positive and negative information, but no evidence that the valence of this information influenced executive control. The evidence of an influence of depressive symptoms on executive control was only observed for one of our depression measures (the sMFQ but not CIS-R), reducing our confidence in the finding. Overall, our findings do not support our hypothesis that adolescents with depressive symptoms would show worse executive control in response to positive than negative information. In our longitudinal investigation, we found no evidence that executive control or sensitivity to positive or negative information was a marker of vulnerability to future depressive symptoms.

### Strengths and limitations

To our knowledge, this is the largest study to examine associations between executive control of positive and negative information and depressive symptoms. The integration of objective neuropsychological measures with epidemiological research methods is a strength of our study. This design should be less prone to selection biases than the previous case-control studies since individuals with and without depressive symptoms are recruited from the same population. Our sample was also considerably larger than both of the prior cohort studies in this area.

Missing data is often a limitation of cohort studies and attrition was high in our sample. We used multiple imputation to minimize biases that might have been introduced by missing data. The wealth of data available in ALSPAC allowed us to identify variables associated with missing data, supporting the plausibility of the missing at random assumption which is a requirement of multiple imputation (Sterne et al., 2009). Results from our imputed sample were consistent with those from our complete case sample, suggesting that missing data did not bias our results. Although our sample was large and population-based, it was recruited from one region in the UK and findings may not generalize to other populations. Adolescents who remained in ALSPAC up to age 18 were also more educated with fewer mental health problems than those who dropped out and this might have introduced selection bias. However, even when attrition is systematic, biases to within-cohort associations in ALSPAC have been found to be minimal, which may explain why our complete case and imputed analyses produced similar findings (Wolke et al., 2009). Still, neuropsychological and neuroimaging studies in nationally representative longitudinal samples are rare and this is an important direction for future research (LeWinn et al., 2017).

The lack of consistent evidence for the MFQ and the CISR also limits our ability to draw conclusions. The CIS-R is, arguably, a more valid measure. It was derived from a standardized diagnostic instrument and although we used the CIS-R as a self-report, this agrees closely with the interviewer administered CIS-R (Lewis et al., 1992).

### Integration with existing findings

Our findings, particularly that there was no influence of valence on executive control, are inconsistent with many of the previous smaller studies (Erickson et al., 2005; Kilford et al., 2015; Kyte et al., 2005; Maalouf et al., 2012; Murphy et al., 1999; Owens et al., 2012). However, findings from these studies are also inconsistent with each other (summarized in Supplementary Table 1). Previous results could be due to selection bias or Type 1 errors. The small samples (Ns < 263) might explain inconsistent findings (Button et al., 2013). There are also several difficulties with the analysis of the affective go/no-go task, especially when sample sizes are small. There are multiple parameters, and multiple approaches to analysis, most of which rely on interaction tests that have low statistical power particularly when the interaction effect is smaller than the main effect (Brookes et al., 2004). This analytical flexibility combined with the small sample sizes increases the probability of a Type I error (Button et al., 2013; Simmons, Nelson, & Simonsohn, 2011).

Our findings are inconsistent with studies reporting generic deficits in executive control in adolescents with depression (Snyder & R., 2013). This could be due to the high cognitive reserve one would expect in an adolescent sample though further studies are required to test this. Executive functions are complex and multifaceted, consisting of lower levels such as concentration and inhibition and higher levels such as reasoning, problem solving, and planning (Diamond & Adele, 2013). The affective go/no-go task assesses lower rather than higher level executive functions. Participants respond very quickly to stimuli they see for a very short period of time (0.3 of a second with under one second to respond). It is possible that the cognitive abnormalities that characterise depression are more dependent on valence when information processing is more deliberative (at the higher level of executive functioning), allowing people more time to apply underlying cognitive schemas or beliefs to information processing. This may happen quickly, and to a certain extent ‘automatically’, but may not be as rapid as the processes assessed by the affective go/no-go. It is also possible that more directly social information is more affected in depression.

## Conclusions

Our evidence does not support the hypothesis that improving executive control of positive and negative information, or even executive control generally, reduces current depressive symptoms or prevents the development of future depressive symptoms in adolescents. Our evidence therefore suggests that improving executive functioning should not be pursued as a strategy for preventing adolescent depression. We conclude that the affective go/no-go task is not a useful neuropsychological task to be pursued in imaging studies of adolescent depression.

## Acknowledgements

We are extremely grateful to all the families who took part in this study, the midwives for their help in recruiting them, and the whole ALSPAC team, which includes interviewers, computer and laboratory technicians, clerical workers, research scientists, volunteers, managers, receptionists and nurses.

## Abbreviations

UK: United Kingdom
ALSPAC: Avon Longitudinal Study of Parents and Children

## References

1. Boyd, A., Golding, J., Macleod, J., Lawlor, D. A., Fraser, A., Henderson, J.,…Davey Smith, G. (2013). Cohort Profile: the ‘children of the 90s’--the index offspring of the Avon Longitudinal Study of Parents and Children. International Journal of Epidemiology, 42(1), 111–127. https://doi.org/10.1093/ije/dys064

2. Brookes, S. T., Whitely, E., Egger, M., Smith, G. D., Mulheran, P. A., & Peters, T. J. (2004). Subgroup analyses in randomized trials: risks of subgroup-specific analyses; Journal of Clinical Epidemiology, 57(3), 229–236. https://doi.org/10.1016/j.jclinepi.2003.08.009

3. Button, K. S., Ioannidis, J. P. A., Mokrysz, C., Nosek, B. A., Flint, J., Robinson, E. S. J., & Munafò, M. R. (2013). Confidence and precision increase with high statistical power. Nature Reviews. Neuroscience, 14(8), 585–586. https://doi.org/10.1038/nrn3475-c4

4. Diamond, A., & Adele, by. (2013). Executive Functions. Annu. Rev. Psychol, 64, 135–168. https://doi.org/10.1146/annurev-psych-113011-143750

5. Elliott, R., Rubinsztein, J. S., Sahakian, B. J., & Dolan, R. J. (2002). The neural basis of mood-congruent processing biases in depression. Archives of General Psychiatry, 59(7), 597–604.

6. Erickson, K., Drevets, W. C., Clark, L., Cannon, D. M., Bain, E. E., Zarate, C. A.,…Sahakian, B. J. (2005). Mood-Congruent Bias in Affective Go/No-Go Performance of Unmedicated Patients With Major Depressive Disorder. American Journal of Psychiatry, 162(11), 2171–2173. https://doi.org/10.1176/appi.ajp.162.11.2171

7. Fraser, A., Macdonald-Wallis, C., Tilling, K., Boyd, A., Golding, J., Davey Smith, G.,…Lawlor, D. A. (2013). Cohort Profile: The Avon Longitudinal Study of Parents and Children: ALSPAC mothers cohort. International Journal of Epidemiology, 42(1), 97–110. https://doi.org/10.1093/ije/dys066

8. Furman, D. J., Hamilton, J. P., & Gotlib, I. H. (2011). Frontostriatal functional connectivity in major depressive disorder. Biology of Mood & Anxiety Disorders, 1(1), 11. https://doi.org/10.1186/2045-5380-1-11

9. Hamilton, J. P., Chen, G., Thomason, M. E., Schwartz, M. E., & Gotlib, I. H. (2011). Investigating neural primacy in Major Depressive Disorder: multivariate Granger causality analysis of resting-state fMRI time-series data. Molecular Psychiatry, 16(7), 763–772.

10. Kilford, E. J., Foulkes, L., Potter, R., Collishaw, S., Thapar, A., & Rice, F. (2015). Affective bias and current, past and future adolescent depression: a familial high risk study. Journal of Affective Disorders, 174, 265–271. https://doi.org/10.1016/j.jad.2014.11.046

11. Kyte, Z. A., Goodyer, I. M., & Sahakian, B. J. (2005). Selected executive skills in adolescents with recent first episode major depression. Journal of Child Psychology and Psychiatry, 46(9), 995–1005. https://doi.org/10.1111/j.1469-7610.2004.00400.x

12. LeWinn, K. Z., Sheridan, M. A., Keyes, K. M., Hamilton, A., & McLaughlin, K. A. (2017). Sample composition alters associations between age and brain structure. Nature Communications, 8(1), 874. https://doi.org/10.1038/s41467-017-00908-7

13. Lewis, G., Pelosi, A. J., Araya, R., & Dunn, G. (1992). Measuring psychiatric disorder in the community: a standardized assessment for use by lay interviewers. Psychological Medicine, 22(02), 465. https://doi.org/10.1017/S0033291700030415

14. Maalouf, F. T., Clark, L., Tavitian, L., Sahakian, B. J., Brent, D., & Phillips, M. L. (2012). Bias to negative emotions: A depression state-dependent marker in adolescent major depressive disorder. Psychiatry Research, 198(1), 28–33. https://doi.org/10.1016/j.psychres.2012.01.030

15. Murphy, F. C., Sahakian, B. J., Rubinsztein, J. S., Michael, A., Rogers, R. D., Robbins, T. W., & Paykel, E. S. (1999). Emotional bias and inhibitory control processes in mania and depression. Psychological Medicine, 29(6), 1307–1321.

16. Nord, C. L., Gray, A., Charpentier, C. J., Robinson, O. J., & Roiser, J. P. (2017). Unreliability of putative fMRI biomarkers during emotional face processing. NeuroImage, 156, 119–127. https://doi.org/10.1016/j.neuroimage.2017.05.024

17. Owens, M., Goodyer, I. M., Wilkinson, P., Bhardwaj, A., Abbott, R., Croudace, T.,…Sahakian, B. J. (2012). 5-HTTLPR and Early Childhood Adversities Moderate Cognitive and Emotional Processing in Adolescence. PLoS ONE, 7(11), e48482. https://doi.org/10.1371/journal.pone.0048482

18. Plichta, M. M., Schwarz, A. J., Grimm, O., Morgen, K., Mier, D., Haddad, L.,…Meyer-Lindenberg, A. (2012). Test–retest reliability of evoked BOLD signals from a cognitive–emotive fMRI test battery. NeuroImage, 60(3), 1746–1758. https://doi.org/10.1016/j.neuroimage.2012.01.129

19. Roiser, J. P., & Sahakian, B. J. (2013). Hot and cold cognition in depression. CNS Spectrums, 18(03), 139–149. https://doi.org/10.1017/S1092852913000072

20. Rothman, K., Greenland, S., & Lash, T. (2013). Modern epidemiology (Third). Philadelphia: Lippincott, Williams and Wilkins.

21. Simmons, J. P., Nelson, L. D., & Simonsohn, U. (2011). False-Positive Psychology. Psychological Science, 22(11), 1359–1366. https://doi.org/10.1177/0956797611417632

22. Snyder, H. R., & R., H. (2013). Major depressive disorder is associated with broad impairments on neuropsychological measures of executive function: A meta-analysis and review. Psychological Bulletin, 139(1), 81–132. https://doi.org/10.1037/a0028727

23. Sterne, J. A. C., White, I. R., Carlin, J. B., Spratt, M., Royston, P., Kenward, M. G.,…Carpenter, J. R. (2009). Multiple imputation for missing data in epidemiological and clinical research: potential and pitfalls. BMJ (Clinical Research Ed.), 338, b2393. https://doi.org/10.1136/BMJ.B2393

24. Tilling, K., Macdonald-Wallis, C., Lawlor, D. A., Hughes, R. A., & Howe, L. D. (2014). Modelling childhood growth using fractional polynomials and linear splines. Annals of Nutrition & Metabolism, 65(2–3), 129–138. https://doi.org/10.1159/000362695

25. Treadway, M. T., & Pizzagalli, D. A. (2014). Imaging the pathophysiology of major depressive disorder - from localist models to circuit-based analysis. Biology of Mood & Anxiety Disorders, 4(1), 5. https://doi.org/10.1186/2045-5380-4-5

26. Turner, N., Joinson, C., Peters, T. J., Wiles, N., & Lewis, G. (2014). Validity of the Short Mood and Feelings Questionnaire in late adolescence. Psychological Assessment, 26(3), 752–762. https://doi.org/10.1037/a0036572

27. Vos, T. (2016). Global, regional, and national incidence, prevalence, and years lived with disability for 310 diseases and injuries, 1990-2015: a systematic analysis for the Global Burden of Disease Study 2015. The Lancet, 388(388), 1545–1602. https://doi.org/10.1016/S0140-6736(16)31678-6

28. Wechsler, D. (n.d.). Wechsler Abbreviated Scale of Intelligence. San Antonio, *TX*: *Psychological Corporation*.

29. Wolke, D., Waylen, A., Samara, M., Steer, C., Goodman, R., Ford, T., & Lamberts, K. (2009). Selective drop-out in longitudinal studies and non-biased prediction of behaviour disorders. British Journal of Psychiatry, 195(03), 249–256. https://doi.org/10.1192/bjp.bp.108.053751

