## Supplementary tables for "Executive control of positive and negative information and adolescent depressive symptoms: cross-sectional and longitudinal associations in a population-based cohort"

| **Study characteristics** | | | | | **Findings** | | |
| --- | --- | --- | --- | --- | --- | --- | --- |
| **Study name (first author)** | **Year of publication** | **Design** | **Sample/setting** | **N** | **Omissions ^a^** | **Commission errors (CE)^b^** | **Reaction times for target words^c^** |
| Murphy | 1999 | Case-control | UK Secondary care patients. Controls recruited by community advertisement | 18 bipolar  28 depressed  22 controls | No evidence | No evidence | Depressed: slower to positive than negative |
| Erickson | 2005 | Case-control | Not clear | 20 depressed  20 controls | Depressed: more positive than negative. Controls: more negative. | No evidence | Depressed: slower to positive than negative |
| Kyte | 2005 | Case-control | UK secondary care adolescents with recent  first episode major depression. Controls from ongoing population-based cohort | 30 depressed  49 controls | No evidence | Depressed: more CE to negative. Controls: more CE to positive. | No evidence |
| Maalouf | 2012 | Case-control | Not clear | 40 depressed (20 acute, 20 remission)  17 controls | No evidence | Depressed: more CE overall | Depressed: faster to negative than positive. Controls and remitted: faster to positive. |
| Owens | 2012 | Cohort | UK sub-sample of larger population-based cohort, selected for high-risk of depression | 238 | Not reported | Neutral & negative CE  later emotional disorder | No evidence |
| Kilford | 2015 | Cohort | UK; children of parents with recurrent depression | 263 | No c-sec evidence. Omissions overall  later depression | Depressed: more positive CE and more positive CE  later depression^d^ | Depressed: faster irrespective of valence |

Supplementary Table 1.

Abbreviations: - associated with; CE - commission errors; c-sec - cross-sectional.

^a^More positive omissions means that more positive target words (correct responses) were missed and same for negative and neutral omissions.

^b^A positive commission error means more commission errors (incorrect responses) when distractor words were positive and same for neutral and negative CE

**^c^**Targets refers to the target word (the correct response)

^d^In shift conditions only.

Supplementary Table 2. Characteristics of the analytic sample (N=2315) compared to the rest of the core ALSPAC sample.

| Characteristic | Complete exposure  (N= 2315) | ALSPAC core sample  (N= 12,135) |
| --- | --- | --- |
| Female offspring, N (%) | 1307 (56.5) | 5437 (46.7) |
| Lower maternal education, O Level or less, N (%) | 1203 (53.7) | 6765 (67.1) |
| Lower maternal social class, N (%) | 347 (18.3) | 1762 (23.1) |
| Offspring depression diagnoses at age 18, CIS-R, N (%) | 138 (6) | 196 (9) |
| Maternal age at birth, mean (SD) | 29.3 (4.5) | 27.7 (5.0) |
| Offspring IQ score, mean (SD) | 92.3 (12.6) | 91.8 (13.2) |
| Offspring MFQ score at 18, mean (SD) | 6.3 (5.2) | 6.8 (5.2) |

Abbreviations: N – number; CIS-R – Clinical Interview Schedule-Revised; MFQ – Mood and Feelings Questionnaire; IQ – Intelligence Quotient; SD – Standard Deviation.
